# The Role of Human Platelet-Rich Plasma to Enhance the Differentiation from Adipose derived Mesenchymal Stem Cells into Cardiomyocyte: An Experimental Study

**DOI:** 10.1101/2020.12.10.420679

**Authors:** I Gde Rurus Suryawan, Andrianto, Arifta Devi Anggaraeni, Arisya Agita, Ricardo Adrian Nugraha

## Abstract

**Background:** Adipose derived mesenchymal stem cells (AMSCs) offer great potential to differentiate into cardiomyocyte. However, the optimal method to maximize the proliferation and differentiation is challenging. Platelet rich plasma (PRP) which contains high levels of diverse growth factors that can stimulate stem cell proliferation and differentiation in the context of cardiac tissue regeneration

**Objective:** To analyze the effect of PRP administration on the AMSCs differentiation into cardiomyocyte and compare to the group without PRP administration.

**Methods:** This study is a true experimental randomized post-test design study. AMSCs were isolated from adipose tissues and cultured until 4 passages. The samples were divided into 3 groups, i.e. negative control (α-MEM), positive control (differentiation medium), and treatment group (PRP). The assessment of GATA-4 marker expression was conducted using flowcytometry on the fifth day and cTnT was conducted using immunocytochemistry on the tenth day to determine the differentiation to cardiomyocyte. Data analysis was conducted using T-test and One-Way ANOVA on normally distributed data determined through Shapiro Wilk test.

**Results:** Flowcytometry on GATA-4 expression revealed significant improvement on PRP group compared to negative and positive controls (67.04 ± 4.49 vs 58.15 ± 1.23 p < 0.05; 67.04 ± 4.49 vs 52.96 ± 2.02 p < 0.05). This was supported by the results of immunocytochemistry on troponin expression which revealed significant improvement on PRP group compared to negative and positive controls (38.13 ± 5.2 vs 10.73 ± 2.39 p < 0.05; 38.13 ± 5.2 vs 26.00 ± 0.4 p < 0.05). This was concordant to the hypothesis which stated that there was an effect of PRP administration on AMSCs differentiation into cardiomyocyte.

**Conclusion:** PRP administration on AMSCs culture significantly improve the differentiation to cardiomyocyte measured by GATA-4 and cTnT expressions.

## INTRODUCTION

Cardiovascular disease remains the leading cause of mortality and morbidity worldwide, causing 17.500.000 deaths in 2012, and contributing to 46.2% of all deaths reported from all over the world in 2014. Data from World Health Organization (WHO) showed that in 2014, 17.9 million people in the world died due to cardiovascular disease, 7.4 million of which died due to coronary heart disease (CHD) (WHO, 2014). Vast majority of patients with CHD will experience into myocardial necrosis that led to contractility dysfunction. Cardiomyocyte has low regenerative capacity; therefore, ischemia or infarct condition will cause pathological cascade causing myocyte hypertrophy, ventricular remodeling, and myocardial fibrosis, all of which will lead to heart failure. Cardiomyocyte constitutes 76% of myocardial structural volume, however the number of cardiomyocytes is only 20-25% of all cells in the heart. Although many measures have been made to cure this condition, poor prognosis and irreversibility of this condition are still widely reported (Minicucci et al., 2011).

Stem cell therapy has demonstrated beneficial effects on several cardiovascular diseases including ischemic heart disease and chronic heart failure. Ischemic cardiomyopathy is a condition that is deemed suitable as a target of stem cell therapy. Mesenchymal Stem Cells (MSCs) is an ideal source of replacement cells because of their potential for self renewal, proliferation and differentiation. MSCs can be abundantly found inside the body, including in bone marrow and adipose tissue. As an addition, MSCs can produce cytokines, chemokines, and growth factors to increase neovascularization, decrease fibrosis in heart, and preserve cardiac function. Therefore, there is possibility that MSCs can be the source of therapeutic cells with the capacity to recover damaged tissues in cardiovascular diseases (Konoplyannikov, 2016).

For a few years, bone marrow-derived MCSs (BMSCs) is a promising therapy strategy in cardiovascular field, however recent studies have shown that adipocyte-derived mesenchymal stem cells (AMCSs) as potential as BMSCs to differentiate into cardiomyocytes under appropriate stimuli (Rodriguez et al., 2005; De Ugarte et al., 2003). Many efforts have been made to facilitate differentiation of MSC to cardiomyocyte, such as administration of 5-azacytidine. However, 5-azatacydine alone is not sufficient to support the differentiation of MSC to cardiomyocyte completely (Wan Safwani et al., 2012).

Platelet rich plasma (PRP) is an autologous plasma which contains concentrated and activated platelet. PRP contains high levels of diverse growth factors that can stimulate stem cell proliferation and differentiation in the context of tissue regeneration. Basic cytokines that can be identified inside the thrombocytes were transforming growth factor–β (TGF-β), platelet-derived growth factor (PDGF), insulin-like growth factor (IGF), fibroblast growth factor (FGF), and vascular endothelial growth factor (VEGF). These cytokines play major role in cell proliferation, chemotaxis, cell differentiation, and angiogenesis (Andia & Abate, 2013). Until recently, the research about the trans-differentiation of MSCs into cardiomyocyte grow rapidly, however there is still no research that explore the effect of PRP on AMSCs differentiation to cardiomyocytes.

## METHODS

### Preparation of AMSCs

AMSCs isolation procedure begins with local anesthesia, followed by minimal surgery of adipose tissue, conducted using standardized method and collected in sterile closed tube, before undergoes medical washing to eliminate any contamination. Before sample is collected, alpha MEM as transport medium is prepared to carry adipose tissue with minimal size of 1 cm^2^ which does not contain any connective tissue or blood clot. Sample obtained is directly put into the transport medium and transported to stem cell laboratory to undergo isolation process. Transportation duration from sample collection location to laboratory should not exceed 1-2 hours.

### Isolation of AMSCs

At the laboratory, adipose tissue is taken out from the transport medium and then washed using PBS solution until red blood cells on the adipose tissue is also eliminated. Adipose tissue is then chopped to small pieces and mixed with collagenase enzyme. It is then poured into bottle containing magnetic stirrer. Tissue inside the bottle is then incubated on hot plate at temperature of 37°C for 30 minutes until the adipose tissue is completely dissolved. After the tissue is dissolved, stopper medium is added and incubated again for 10 minutes until the solution become homogenous. The tissue was then poured on 50 mL conical with sterile gauze on top of it so that undissolved tissue remnants can be separated. The solution is then centrifuged with speed of 3000 rpm for 4 minutes until pellet is formed. Pellet is then resuspended with alpha MEM medium until it becomes homogenous solution. After that, it is planted on 10 cm petri dish and incubated inside CO_2_ incubator for 24 hours until the cells stick to the base of petri dish. Medium is then replaced every 2 days until the cells form colonies and grow until it reaches 80% confluency.

### Identification of AMSCs phenotype from Adipose Tissue

Isolated MSCs from adipose tissue need to undergo characterization using CD105 which is specific marker for MSCs. After that, MSCs that has been labelled is assessed using flow cytometry and immunocytochemistry. Positive test result towards CD105 is indicated by fluorescence on MSCs membrane surface. CD45 and CD34 which are specific markers for hemopoietic stem cells need to be used to make sure that the isolate from adipose tissue is purely MSCs, shown by no fluorescence on MSCs membrane surface (negative result).

### Culture of AMSCs

MSCs that have grown into colony could be multiplied into needed dose for clinical application. Cells which have formed monolayer with 80% confluency should be rejuvenated by doing passage. Passage is done by removing medium from petri dish and washing the monolayer with PBS solution. After that, triple express enzyme is added and incubated for 5 minutes until monolayer is detached from the base of the petri dish. After detachment of monolayer, stopper medium is added and resuspension is done until single cell is formed. The solution containing single cell is poured into conical tube and centrifuged until pellet is formed. Pellet is then cultured in alpha MEM medium and differentiate into cardiomyocytes, after that resuspension is done until homogenous solution is formed and planted in new petri dish.

### Preparation of PRP

Blood that has been obtained from the patient is poured into 50 mL conical and centrifuged with speed of 3000 rpm for 10 minutes until two layers are formed, which are plasma on top layer and blood cells on bottom layer. The top layer is collected slowly and poured into a new 50 mL conical tube and centrifuged again for 10 minutes with the speed of 3000 rpm until white pellet is formed on the bottom. The upper two-thirds is removed, and the lower one-third is resuspended again until the solution becomes homogenous. The solution is then filtered using 0.22 micro and PRP sample is ready for treatment.

The result of AMSCs culture in this fourth step will be divided into three groups. First group is negative control group, which is AMSCs culture group on alpha mem medium as standard medium. Second group is positive control group, which is AMSCs culture on differentiation medium without PRP administration. The third group is treatment group, which is AMSCs culture group on differentiation medium with PRP administration.

### Cardiomyocytes Differentiation Assay

Cells that have been multiplied and reached 80% confluency can be subculture on 24 well plate. Seeding of those MSCs is done for 4 consecutive days, the medium is regularly changed every day in certain order. On day 1, medium is changed into alpha MEM, on day 2 medium is changed into A differentiation medium, on day 3 medium is changed into B differentiation medium, and on day 4 medium is changed into maintenance medium. After cells grow confluently, on fifth day cell is ready to be harvested and assessed using flowcytometry.

Cells that have been multiplied and reached 80% confluency is ready to be subculture on well 24 that has been given coverslip. Immersion of the coverslip needs to be done by adding alcohol 70% for 5 minutes and then washed using PBS solution and lastly culture medium is added before MSCs seeding. Seeding of those MSCs is done for 4 consecutive days, the medium is regularly changed every day in certain order. On day 1, medium is changed into alpha MEM, on day 2 medium is changed into A differentiation medium, on day 3 medium is changed into B differentiation medium, and on day 4 medium is changed into maintenance medium. After cells grow confluently, on fifth day cell is ready to be harvested and assessed using immunocytochemistry by removing cell medium in 24 well plate, and then washed using PBS and fixated with formaldehyde 4% for two hours. Cell is ready to be assessed using immunocytochemistry staining on tenth day.

### Study Design

This study is laboratory experimental study (in vitro study) by administrating plasma rich platelet (PRP) on AMSCs culture. The objective is to assess the effect of PRP administration towards AMSCs ability to differentiate into cardiomyocytes. This study is a true experimental randomized study, with “post-test only control group design” approach (Figure 1). Experimental study uses three principles; randomization, replication and control. In this study, these three principals were applied. Randomization was done before the samples were divided into three groups. Replication principle increased accuracy of the result with minimal variation. In replication principle of statistics, it must be done for minimum three times so that calculation of standard deviation can be done. This was also in accordance with triplicate calculation principle. After that, negative and positive control were made. Randomization technique of samples in this study was based on random result of AMSCs samples with good qualities according to criteria made by Stem Cell Team of Biomaterial Centre and Tissue Bank of Dr. Soetomo General Academic Hospital, Surabaya. This study was conducted for three months (November 2019-January 2020),

**Figure 1.**
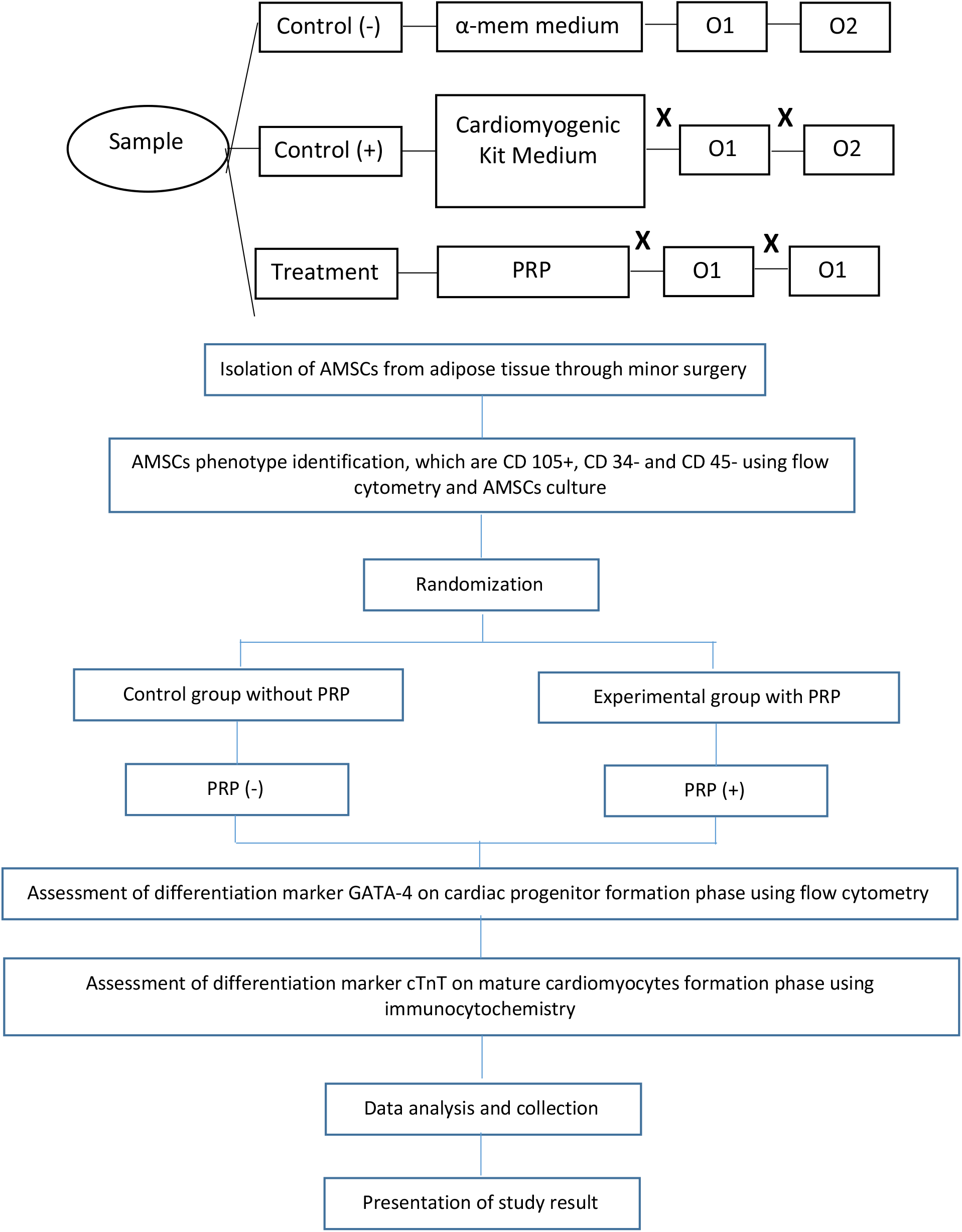
Research Flowchart

### Data Analysis

Data collected in this study will undergo several processes including coding, entry, cleaning, and editing. Data collected in this study is primary data, through flow cytometry and inverted microscope. Obtained data will be inputted and processed using statistic software SPSS version 25.0. Data will be presented in form of mean / median and value range and will be presented in the form of table or graph. Then data will undergo inferential analysis, using one-way ANOVA if the distribution is normal, or Kruskal Wallis if the distribution is not normal. To analyze the difference between two groups (post hoc), T test will be used if the distribution is normal, or Mann Whitney test if the distribution is not normal. Significance cut off value is α = 0.05. In this study, because the sample size is < 30, Shapiro-Wilk test will be used (Figure 1).

## RESULTS

### Isolation and Culture of AMSCs

AMSCs was isolated from adipose tissue, which was obtained using minimal invasive method. This isolation procedure was in concordance with the standard from stem cell laboratory inBiomaterial Centre and Tissue Bank of Dr. Soetomo General Academic Hospital. After being isolated, AMSCs were cultured and expanded until fourth passage with the objective to maintain stemness characteristic of AMSCs and prevent further cell differentiation. Majority of the cell is spindle-like shaped if seen under the microscope with 100 times magnification. Below are pictures of AMSCs cultures microscopically, starting from isolation (passage 0) until passage 4 (Figure 2).

**Figure 2.**
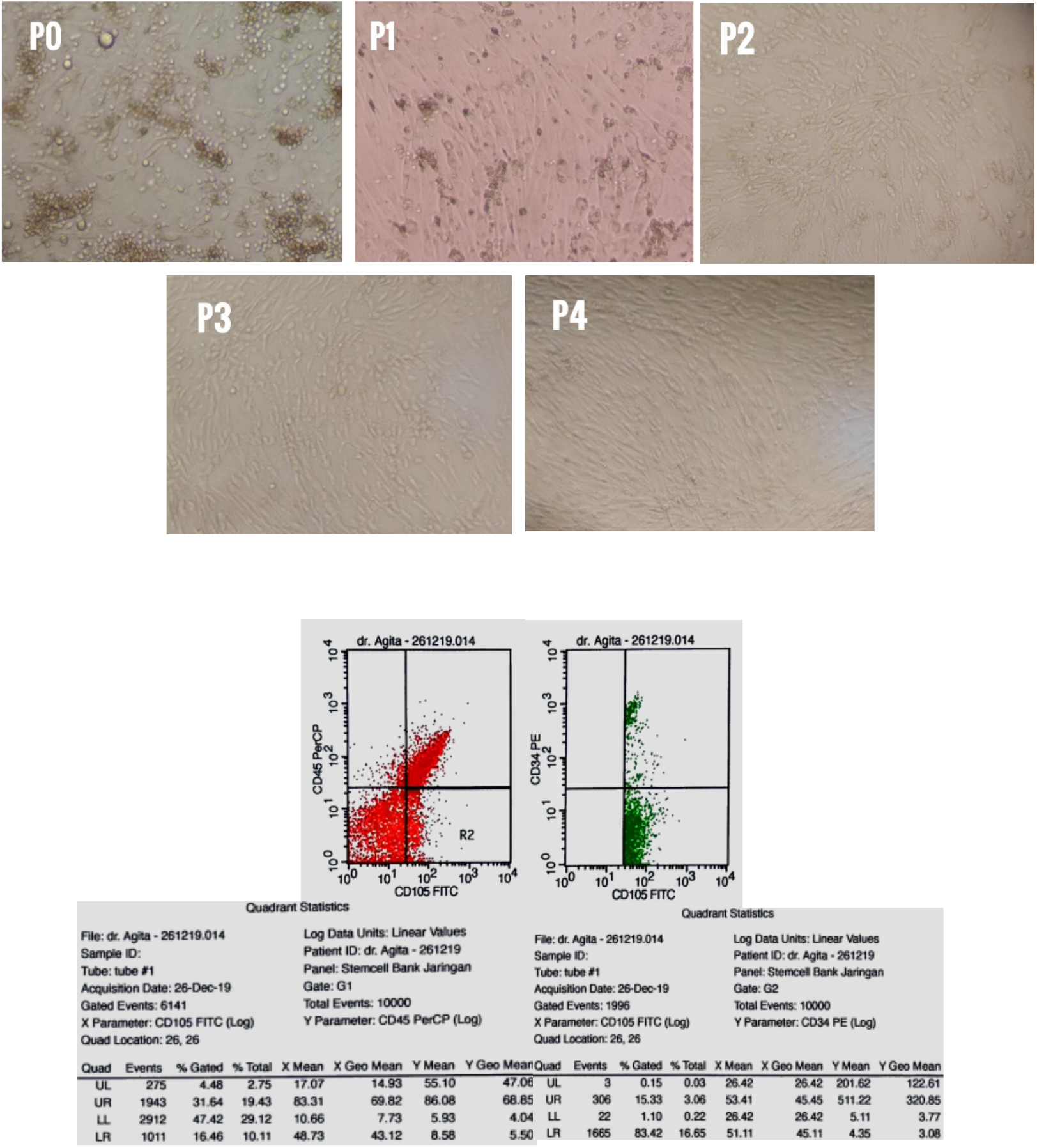
AMSCs Isolation, culture, and characteristics on flow cytometry

### AMSCs Phenotype Characterization

The minimum criteria for MSCs characterization by The International Society for Cellular Therapy (ISCT) is that most of them show positive expression of CD73, CD90, and CD105, and negative expression of CD11b or CD14, CD19, CD34, CD45, and HLA-DR. In our study, AMSCs obtained from adipose tissue will undergo MSCs characterization on fourth passage culture. AMSCs culture will show positive expression of CD105^+^ and negative expression of CD34^−^ and CD45^−^ with flow cytometry. Phenotype identification morphologically using inverted microscope would show adherent cell (Figure 2).

### Preparation, isolation, and PRP administration on treatment group

Blood that had been aspirated from the patient was poured into 15 mL conical and centrifuged with speed of 3000 rpm for 10 minutes until two layers are formed, which were plasma on top layer and blood cells on bottom layer. The top layer was collected slowly, poured into a new 15 mL conical tube and centrifuged again for 10 minutes with the speed of 3000 rpm until white pellet was formed on the bottom. The upper two-thirds were removed, and the lower one-third was resuspended again until the solution became homogenous. The solution was then filtered using 0.22 micro and PRP sample was ready for treatment. PRP dose which was given to each group was 5 × 10^9^/L (Figure 3).

**Figure 3.**
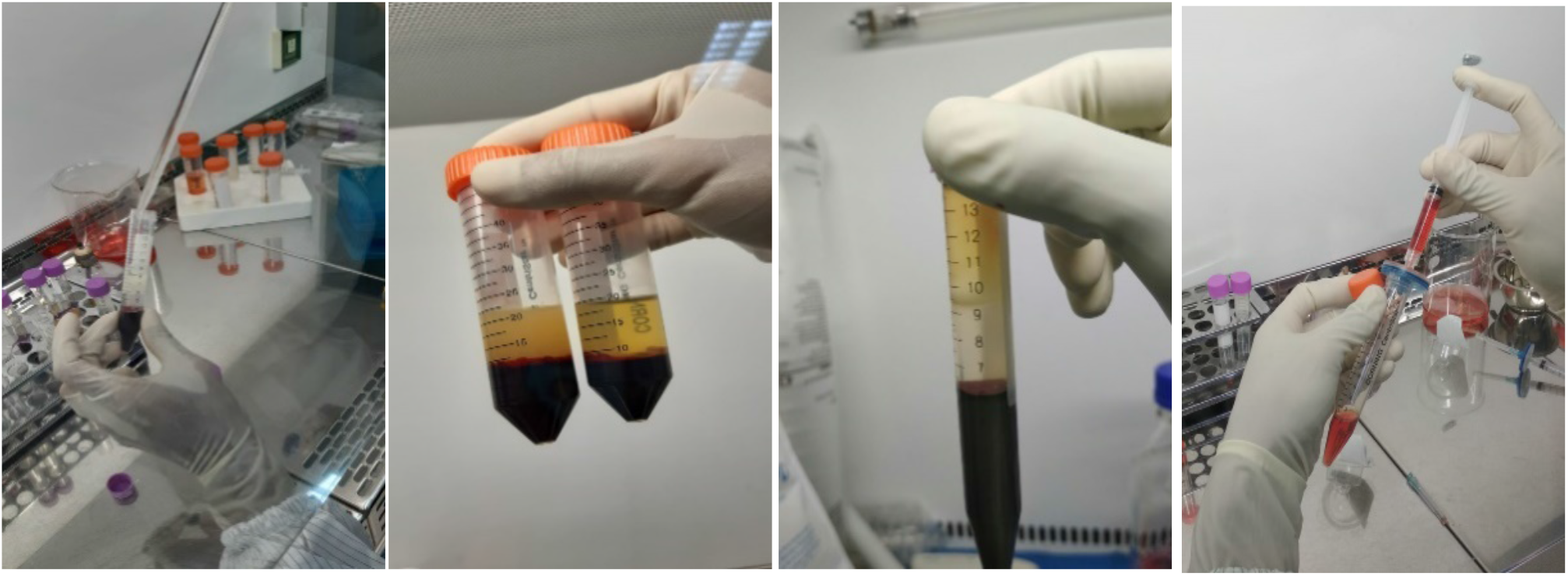
PRP formation steps

### Assessment of cardiomyocytes differentiation marker GATA-4 expression using flow cytometry

Differentiation into cardiomyocytes happened through several cell development and maturation steps, indicated by expression of specific transcription factors. According to sequential diagram of cardiomyocytes development steps from pluripotent cells with typical marker expressed on each step by Rajala et al, 2018, GATA-4 is one specific cardiac transcription factor that indicates development of cardiomyocyte on cardiac progenitor formation phase. Assessment of GATA-4 in this cardiomyocyte differentiation step can be done on day 5-7, in this study assessment of GATA-4 using flow cytometry was done on fifth day because at that time AMSCs had already looked confluent. Flow cytometry is a technique used to analyze cells types in cell population. To assess the difference between those three groups, distribution test was done to assess the normality of data distribution. Normality test was done using Shapiro Wilk test. Data distribution is considered normal if P value > 0.05, in this study the data distribution was normal with *p* value = 0.739 (Figure 4).

**Figure 4.**
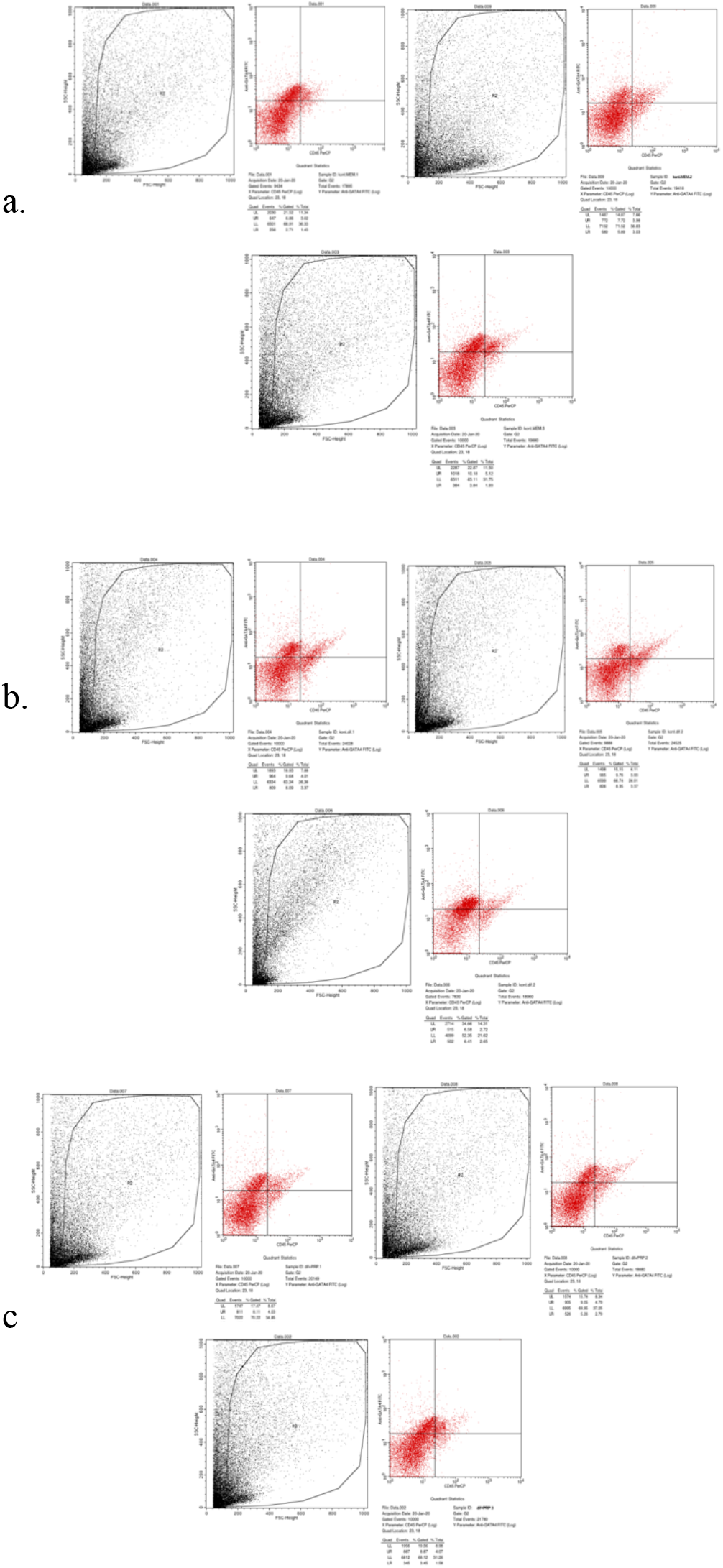
Flow cytometry from: a) alpha MEM medium group; b) differentiation medium group; c) treatment group (with PRP administration on differentiation medium)

From flow cytometry assessment of three groups, mean ratio between expression of GATA-4 (right upper quadrant) and no expression of GATA-4 (right lower quadrant) in AMSCs group was 58.1467, with standard deviation of 1.23 in negative control group (alpha-MEM), 52.9633 with standard deviation of 2.02 in positive control group (differentiation medium), and 67.04 with standard deviation of 4.49 in treatment group (differentiation medium + PRP). These data showed that the mean ratio of treatment group was superior (higher) than negative control group and positive control group (Figure 5).

**Figure 5.**
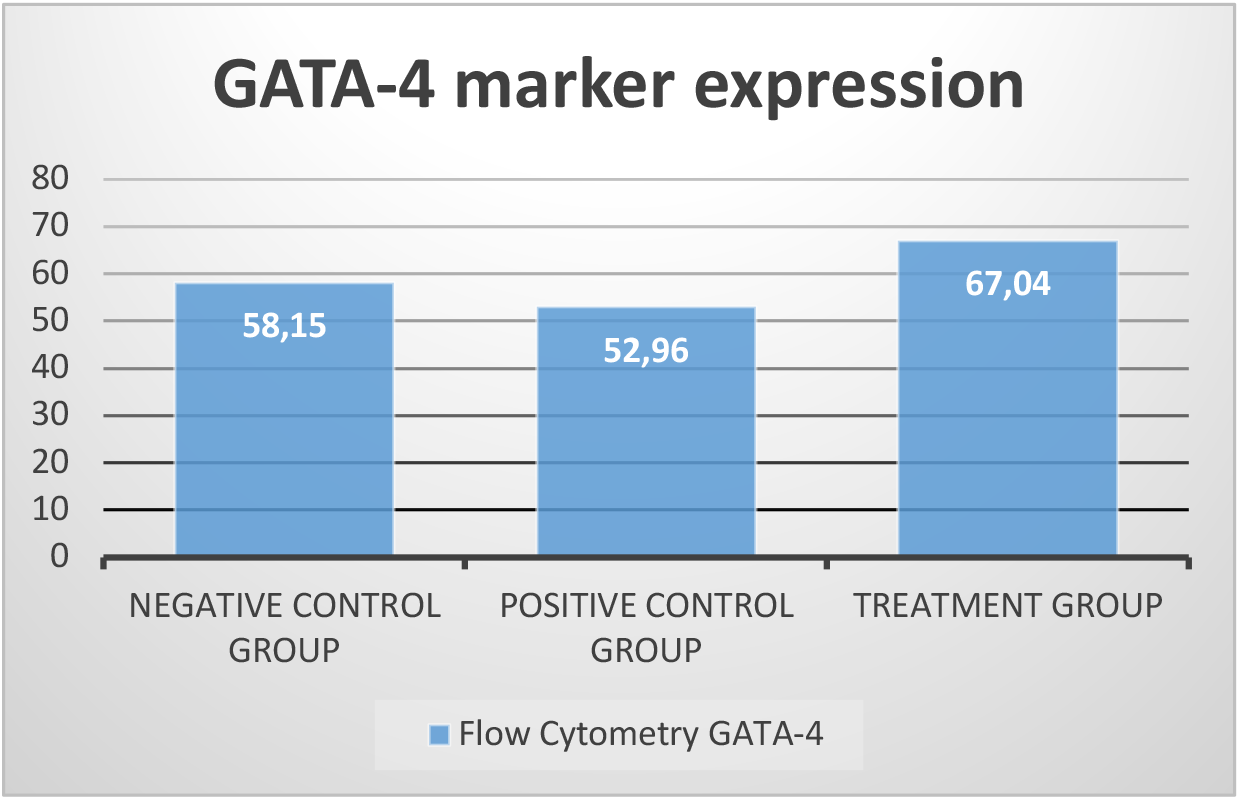
Bar chart showing mean ratio of GATA-4 expression (right upper quadrant) and no expression of GATA-4 (right lower quadrant) in AMSCs group.

Due to normal distribution of the data, ANOVA test was done to assess the difference between three sample groups, and the result was *p* = 0.003. This result showed that there was significant difference in ratio of GATA-4 (right upper quadrant) and no expression of GATA-4 (right lower quadrant) between those three groups. T-test was done to assess the difference between two groups. The result showed that there was significant difference between negative control group (alpha-MEM) compared with treatment group (differentiation medium + PRP) and positive control group (differentiation medium) with p < 0.05. However, no significant difference was found between negative control group (alpha-MEM) and positive control group (differentiation medium) with *p* = 0.074 (Table 1).

**Table 1.** T-test between negative control group, positive control group, and treatment group using flow cytometry

| T-test parametric test | Mean difference | Significance cut-off |
| --- | --- | --- |
| α-MEM to differentiation medium | 5.18333 | 0.074 |
| α-MEM to PRP | 8.89333 | 0.010 |
| Differentiation medium to PRP | 14.07667 | 0.001 |

### Assessment of cardiomyocytes differentiation marker troponin expression using immunocytochemistry

Troponin (cTnT) is final marker of cardiomyocytes which can be detected starting from tenth day and reach its peak on 14th day. On tenth day, the three groups were assessed for cTnT marker expression using immunocytochemistry. From previous studies, it was found that the ideal timing to assess AMSCs differentiation into cardiomyocytes is on the tenth day. On immunocytochemistry testing, cardiomyocytes can be identified morphologically which is visualized by cytoplasm staining with DAB chromogen and nuclear staining with Meyer hematoxylin. Cardiomyocytes would be described to be more prominent compared with other cells, with prominent blue nuclear and prominent brown cytoplasm in 400x magnification. On those three groups, the total amount of cardiomyocytes was counted from each microscope field of view. To assess the difference between those three groups, distribution test was first done to know the normality of data distribution. Because the sample size was < 30, normality test used was Shapiro Wilk. Data distribution was said to be normal if P value > 0.05, in this study the data distribution was normal with *p* = 0.537 (Figure 6).

**Figure 6.**
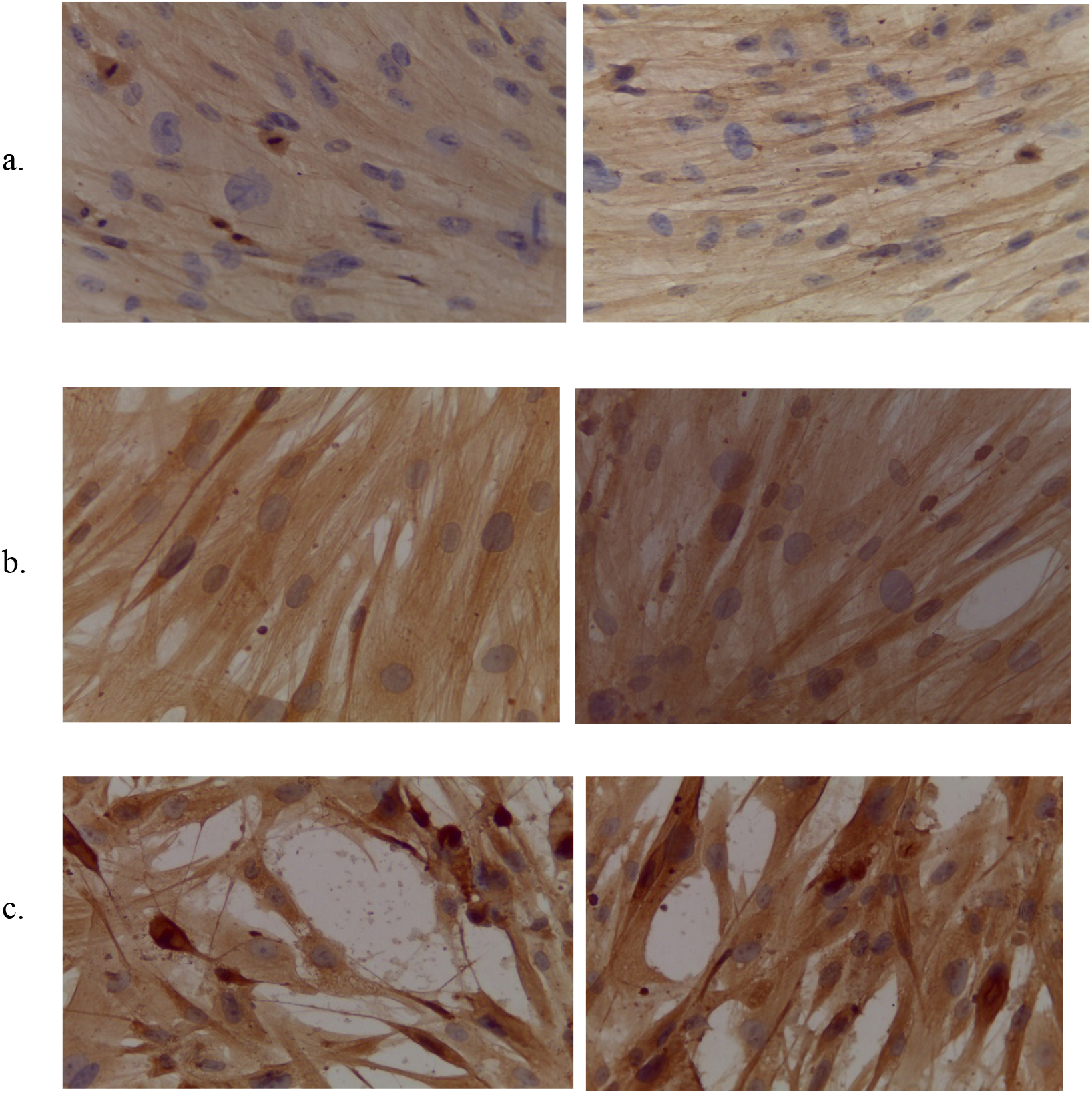
Immunocytochemistry from: a) alpha MEM medium group; b) differentiation medium group; c) treatment group (with PRP administration on differentiation medium)

From immunocytochemistry assessment of those three groups, the mean value was 10.73 with standard deviation of 2.39 in negative control group (alpha-MEM), 26.0 with standard deviation of 0.4 in positive control group (differentiation medium) and 38.13 with standard deviation of 5.2 in treatment group (differentiation medium + PRP). This result showed that treatment group was superior to negative and positive control group by looking at the mean value (Figure 7).

**Figure 7.**
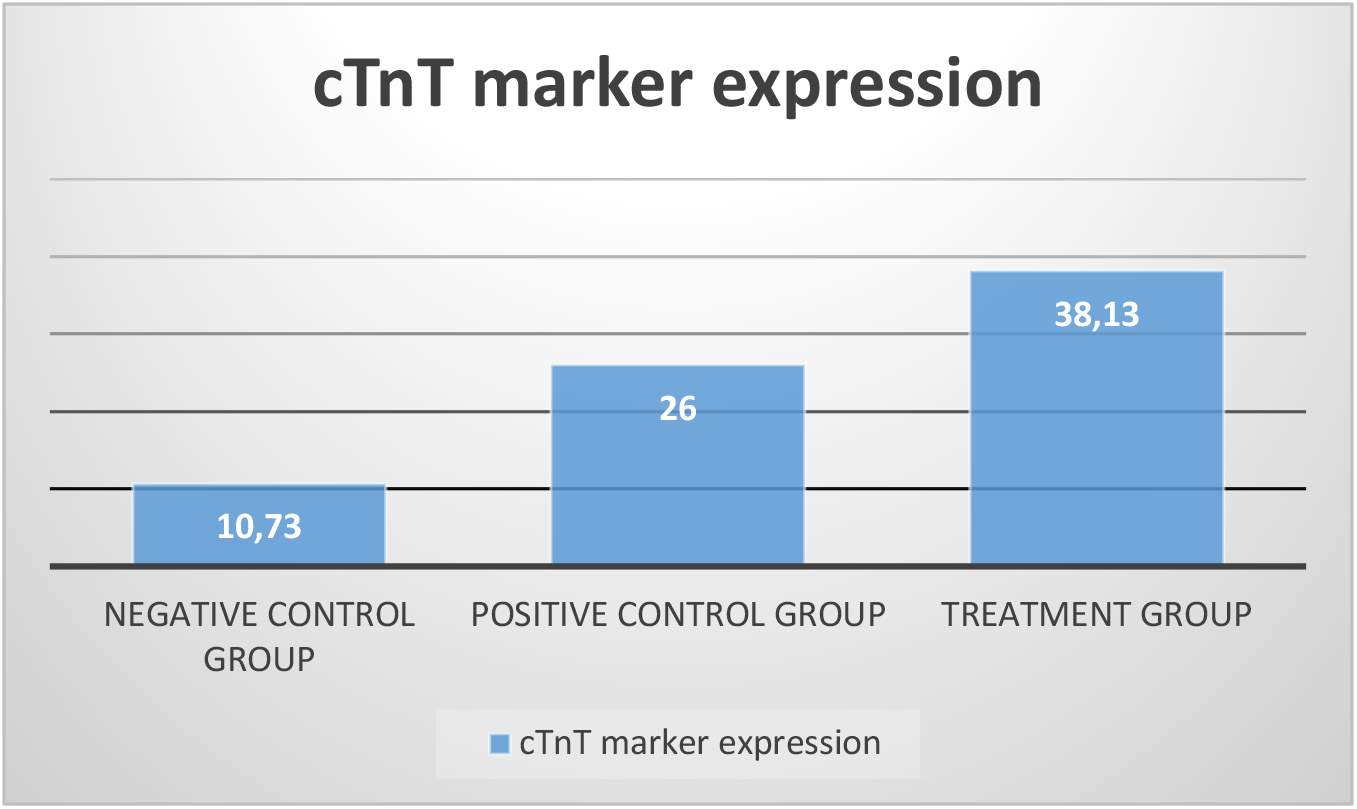
Bar diagram showing mean of cardiomyocytes through expression of troponin

Because the data distribution was normal, ANOVA test was done to assess the difference between those three groups, the result was p = 0.0001. This implied that there was significant difference in mean value of cardiomyocytes, expressed by troponin, between those three groups. T-test was also done to assess difference between two groups. There was significant difference between negative control group (alpha-MEM) and treatment group (differentiation medium + PRP), negative control group (alpha-MEM) and positive control group (differentiation medium), and positive control group (differentiation medium) and treatment group (differentiation medium + PRP) with p < 0.05 (Table 2).

**Table 2.** T-test between negative control group, positive control group and treatment group using immunocytochemistry

| **T-test parametric test** | **Mean difference** | **Significance cut-off** |
| --- | --- | --- |
| α-MEM and differentiation medium | 15.27 | 0.001 |
| α-MEM and PRP | 27.4 | 0.000 |
| Differentiation medium and PRP | 12.13 | 0.004 |

## DISCUSSION

AMSCs have the same therapeutic potential as BMSCs, which are mesenchymal stem cell with multipotent characteristic, and can differentiate into several cell types such as osteocyte, chondrocyte, adipocyte, cardiomyocyte, and fibroblast. Minimal criteria to define MSCs culture according to ISCT 2006 are; (a) adherence to plastic in standard culture media, (b) positive specific surface antigen (Ag) expression, which are CD73, CD90 and CD105; and negative expression of CD11b or CD14, CD19 or CD79α, CD45, CD34 and HLA-DR, and (c) multipotent differentiation potential, i.e., can differentiate into adipocyte, chondrocyte, osteoblast using staining in cell culture in vitro. This study used phenotype characteristic, which are expression of CD105+, CD 34−, and CD 45−, on AMSC passage-4 culture with flow cytometry technique. This is in accordance with minimal criteria for MSCs characterization defined by ISCT (Raposio, 2016).

Patient with post myocardial infarction and heart failure will develop ischemic cardiomyopathy if not given adjuvant therapy. Some of these patients are said to be refractory to pharmacological therapy. In this case, stem cell therapy is expected to be advanced therapy for treatment of ischemic cardiomyopathy. Platelet functionally has major role in tissue regeneration. However, until now, there is still no study explaining the role of PRP on AMSCs differentiation into cardiomyocytes. PRP potential role in increasing healing of bone, muscle, ligament, and tendon tissue has been applied in almost all orthopedic sub specialization (Weibrich G., et al, 2004). In plastic surgery, PRP is also said to be the source of protein for wound healing and tissue regeneration. This study aimed to assess the effect of PRP administration toward AMSCs differentiation into cardiomyocytes in vitro. PRP, which is rich in growth factor, was expected to increase AMSCs differentiation into cardiomyocytes.

Marx and Dohan have made classification about the role of PRP concentration and its efficacy, however this classification has not been used widely. This study used DEPA standard classification, in which the dose of platelet injected is measured in billion or million and categorized as follow: very high dose > 5 billion, high dose 3-5 billion, intermediate dose 1-3 billion, and low dose < 1 billion. Considering the information available in four publications, platelet dose can be counted with initial platelet concentration of 200 × 10^9^/L. It is said that PRP optimal dose is > 5 billion, where red blood cells contamination is very low. That very high dose can be achieved by obtaining minimum of 30 cc of venous blood. This study used that very high dose to get optimal study result (Magalon et al. 2016).

After myocardial infarction, the heart will undergo great inflammatory process for 3-4 days (in rats) which involve inflammatory gene regulation and immune cells infiltration into interstitial space of myocardium, to remove damaged cardiomyocytes. This acute inflammation period is fundamental to tissue homeostasis. Heart fibroblast was traditionally regarded as an important component to give structural and functional integrity to myocardium. Many activation fibroblast mediators identified, including serotonin, TGF-β1 and PDGF, all of which are contained in high amount in platelet granules, stimulating fibroblast expansion and trans-differentiation into myofibroblast (Travers JG, et al., 2016).

After coronary thrombosis, activated platelet will release granules component and microparticles which regulate: 1) extravasation and accumulation of inflammatory cells inside myocardium after myocardial infarction, 2) influencing immuno-active response of leukocytes, especially neutrophile and monocyte/macrophage (M1) and allowing M2 activity for tissue regeneration, 3) fibroblast activation and conversion into myofibroblast to promote extracellular matrix (ECM) synthesis, 4) proliferation, mobilization, and differentiation of progenitor cell (cardiac progenitor), 5) differentiation of progenitor cell into cardiomyocyte to improve cardiac remodeling and repairment, and 6) cardiomyocytes are sensitive to platelet molecules release, which can increase inotropic activity of cardiomyocytes and give anti apoptotic signal (TGF-β1) (Walsh Tony, Poole Alastair, 2018).

In addition, platelet alpha-granule can also produce chemokine stromal cell-derived factor-1α which regulate progenitor cell differentiation into cardiomyocyte. Platelet can also release several spectrums of miRNA (including miRNA-199 and miRNA-29) which can influence cardiomyocyte reentry cycle (Eulalio A, et al., 2012). Some of the theories support the result of this study; it is said that PRP can increase AMSCs differentiation into cardiomyocytes, and there was significant difference in PRP administration group compared with negative control group (alpha-MEM) and positive control group (differentiation medium without PRP administration) assessed by GATA-4 marker expression (67.04 ± 4.49 vs 58.15 ± 1.23 *p*<0.05; 67.04 ± 4.49 vs 52.96 ± 2.02 *p*<0.05) on cardiac progenitor formation phase and troponin marker expression (38.13 ± 5.2 vs 10.73 ± 2.39 *p*<0.05; 38.13 ± 5.2 vs 26.00 ± 0.4 *p*<0.05) on cardiomyocytes formation phase. In conclusion, platelet plays important role in cell repair through released biomolecules (Walsh Tony, Poole Alastair, 2018).

## CONCLUSION

In summary, our study demonstrated that PRP can induce the differentiation of AMSCs into cardiomyocyte by releasing growth factors including PDGF, FGF, IGF and TGF-β. There was difference on AMSCs differentiation into cardiomyocyte with PRP administration compared to α-MEM medium and Cardiomyogenic Kit medium. This was documented by expression GATA-4 on the fifth day and cTnT on the tenth day. Thus, PRP which is simple and lower cost to prepare, greatly improves the differentiation of AMSCs into cardiomyocyte by in vitro study. Further study is required with more comprehensive differentiation markers, starting from cardiac mesoderm formation to mature cardiomyocyte. Flow cytometry and immunocytochemistry assessment is needed in every differentiation step to reduce positive bias of study result. Microscopic examination with inter and intra observer is also needed to reduce positive bias resulting in more accurate interpretation.

## AVAILABILITY OF DATA AND MATERIAL

The datasets generated during and/or analyzed during the current study are not publicly available due to protecting participant confidentiality but are available from the corresponding author on reasonable request.

## ACKNOWLEDGMENT

The authors thank to Stem Cell Team of Biomaterial Centre and Tissue Bank of Dr. Soetomo General Academic Hospital, Surabaya.

## FINANCIAL SUPPORT AND SPONSORSHIP

Nil.

## CONFLICT OF INTEREST

The authors declare that they have no competing interests.

## Notes

http://repository.unair.ac.id/96614/

### Competing Interest Statement

The authors have declared no competing interest.

### Summary of Updates

Minor correction to provide institutional email addresses

